# A fractal pattern of hierarchical genetic population structure in mixed stocks across fish segregated by dams revealed by genomic resources for curimba *Prochilodus lineatus*

**DOI:** 10.1101/2025.03.14.643328

**Authors:** Gabriel M. Yazbeck, Gésica Aparecida Santana Nascimento, Leiliane Campos Carvalho, Raíssa Cristina Dias Graciano, Rosiane de Paula Santos, Rafael Sachetto Oliveira

## Abstract

Genomic resources, new microsatellite markers and a novel observation of a fractal pattern of the genetic population structure are presented for curimba *Prochilodus lineatus*, a suitable model organism for freshwater migratory species in South America. Our main goals were to investigate the presence of mixed fish stocks, the effects of isolation by dams and broodstocking over the genetic diversity for this species. We focused on the upper segment of the extremely fragmented Grande River, MG, Brazil, from the megadiverse continental La Plata Basin. Two main co-occurring genetically admixed stocks were identified by maximum likelihood estimation and iteratively unfolded with two further steps, through a hierarchical analysis of genetic structure. It consistently unveiled two newly defined internal groups, from the previously resolved clusters (2^3^). We suggest this stratified pattern of mixed stocks from the main river channel may be attributed to the existence of different, well defined breeding grounds, nested within differential scales throughout this basin, arranged according to fractal geometry (1^st^ order river, 2^nd^ and 3^rd^ order tributaries, and so on). This would imply in homing behaviour for this species. Our work also provides massive molecular data for this fish, including its first draft genome assembly and a valuable panel for the rapid and low-cost development of new microsatellite DNA markers. These findings will likely benefit researchers and environmental managers in tackling challenging issues regarding the conservation of Neotropical migratory fishes, as they support the hypothesis of the isolating effects of dams and broodstocking over genetic variability. More importantly, the results presented here point to the potential role of tributaries of different orders in harbouring genetically diverse populations of migratory freshwater fishes.

## Introduction

The curimba *Prochilodus lineatus* Valenciennes 1836 (Characiformes: Prochilodontidae), also known as sabálo, grumatá and other common names, is one of the most important freshwater migratory species for inland fisheries in South America. It corresponds to >50% of the fish biomass from the La Plata Basin (Bowen, 2022a,b), a megadiverse hydrological system (Cassemiro *et al*., 2023), reaching Argentine, Bolivia, Brazil, Paraguay and Uruguay. This major Neotropical basin is heavily impacted by anthropogenic activity (Reis *et al*., 2016). *Prochilodus lineatus* also naturally occurs in the Paraíba do Sul Basin (Brazil), and has been recorded as an invasive species in China (Endruweit, 2014) and Vietnam (Kalous *et al*., 2012).

*Prochilodus lineatus* has a conspicuous specialized inverted funnel mouth, an adaptation from its detritivorous habits (Bowen, 2022b) and, due to its abundance and popularity, it is an emblematic example of the Neotropical potamodromous (*i*.*e*. freshwater migratory) life history. Its reproductive migrations are triggered by environmental cues and its fish runs are regionally denominated piracema, a term used to refer to both, the migratory process, and the fish species which partake in it. It typically occurs from around the early summer, until around the season’s end (Carolsfeld *et al*., 2003). Thus, this process is potentially sensible to ongoing climate change (Alix *et al*., 2020).

Piracema fishes naturally spawn upon reaching upstream breeding grounds (Ribolli *et al*., 2020). Larvae and juveniles then passively drift downstream and reach nursery and feeding floodplains, where they grow and mature (Petry *et al*., 2003). Adults spawn several consecutive years during its live cycle (Carolsfeld *et al*., 2003).

Species from the *Prochilodus* genus were among the first Neotropical migratory fishes to have its *ex situ* spawning mastered. This was motivated by the intensive exploitation of hydroelectricity generation and its impact over migratory species, aiming broodstock release operations, intended to counteract dwindling fisheries around regions affected by dams (Saraiva and Pompeu, 2016).

These stocking practices have been largely abandoned in some relevant places for hydroelectric production (*e*.*g*. the Brazilian state of Minas Gerais), during the last decade, in part because of general relaxation of environmental protection oriented regulations (Zellhuber, 2016), although a rigorous, holistic (*i*.*e*. ecological, genetic and anthropological) assessment of the activity’s potential benefits or pitfalls is still lacking. Given its status as the main targeted fish for broodstocking, in the last 50 years or so, *P. lineatus* is a natural candidate as a model species for gauging the impacts of dams and the effectiveness of possible mitigation measures, regarding piracema fishes.

Molecular markers have been applied to access the influence of dams over Neotropical migratory fishes showing, for example, the existence of different stocks across fragmented basins (*e*.*g*. Oliveira-Farias *et al*., 2022), or the lack of differentiation following recent isolation (*e*.*g*. García-Castro and Márquez, *2024). It was also used to show the absence of regional genetic structure (e*.*g*. Sivasundar *et al*., 2001), or to approach fish stocks delimitation (Collins *et al*., 2013). DNA markers were also used to reveal life history traits of silurid catfish species, showing evidence for the occurrence of homing breeding behaviour (*i*.*e*. philopatric spawning), in the Amazon (Batista and Alves-Gomes, 2006; Carvajal-Vallejos *et al*., 2014) and the La Plata Basin (Pereira *et al*., 2009). Other questions worth a deeper assessment in South America include the efficiency of transposing mechanisms and practices for sustaining gene flow (*e*.*g*. Wilkes *et al*., 2019); if broodstocking are inducing genetic effects (*e*.*g*. bottlenecks, domestication) from hatchery spawning (*e*.*g*. Howe *et al*., 2024); if released fish lead to stocks introgression (*e*.*g*. Chang *et al*., 2022), or jeopardize local adaptations (*e*.*g*. Shedd *et al*., 2022), and so on.

Piracema fishes under the impacts of hydroelectric power production, thus, may benefit from objective genetic information raised through molecular markers such as microsatellites, which are amenable to simple genotyping routines for hatchery or wild fish monitoring operations (Wenne, 2023). The popularization of next-generation sequencing (NGS) techniques have made the characterization of new DNA markers increasingly accessible and straightforward (Yazbeck *et al*., 2024).

Since the seminal work of Smouse *et al*. (1990), scientists and managers have applied the statistical inference of genetic structure (the form in which genetic polymorphisms are distributed), to characterize mixed stocks from fish samples, through DNA markers (Hallerman, 2003; Valenzuela-Quiñonez, 2016). Migratory runs potentially place individuals from different, well defined, random breeding populations (demes) together in the same local mixed fish stock, along commonly shared migratory routes (Collins *et al*., 2013). Mixed fish stocks can follow a genetically mixed population model, characterized by each individual’s genome originating from a single ancestral population source (genetic cluster), out of a K number of alternatives, where these demes do not share breeding grounds and events. Alternatively, mixed fish stocks might be portrayed by an admixture model. It considers different individual proportions of genomic heritage, out of several K ancestral genetic clusters, where previous generations have potentially engaged in common reproductive events (Pritchard *et al*., 2000; Wang, 2022). Admixture, thus, provide a convenient framework for the problem of unravelling the genetic structure of piracema fishes, under the impact of dams and decades of stocking operations, largely conducted without planned considerations towards genetic issues.

Genetic investigations applying microsatellites and admixture models in *P. lineatus* were first presented by Rueda *et al*. (2013), unveiling distinct seasonal segregation of stocks from the Lower Uruguay River. They have concluded this species should be deemed as exhibiting mixed stocks, for fisheries management purposes. Perini *et al*. (2021) described a genetic source-sink metapopulation dynamics involving the Pardo River and the Lower Grande River (Upper Paraná Basin). Both, Ferreira *et al*. (2017) and Rosa *et al*. (2022) also combined mitochondrial sequences along microsatellites. The former studied fish from the Middle and Upper Paraná River Basin and showed no evidence for homing in this species. The latter revealed temporal genetic structuring of a location of the Mogi-Guaçu River (Pardo River Basin). Ferreira *et al*. (2023) also showed temporal structuring and mixed stocks of *P. lineatus* from the Paranapanema River Basin, a major tributary of the Upper Paraná River.

In general, all these studies relate to lower regions of the same intensely dam fragmented system and discussed the importance of free-flowing tributaries for harbouring spawning groups and events in *P. lineatus*. All showed high levels of genetic variability and frequently reported on the absence of a single panmitic population associated with distinct localities. Instead, these studies point to local or temporal genetic substructuring and tend to resolve two or three genetic clusters, according to Bayesian admixture analysis. To date, except for observations provided by Yazbeck and Kalapothakis (2007), there are no genetic population structure analysis for *P. lineatus* from the Upper Grande River, the final and highest region from the Upper Paraná Basin and, therefore, the uppermost terminal region from the vast Upper La Plata River Basin’s oriental flank.

The Grande River springs from an altitude of almost 2,000 m in Minas Gerais and runs through 1,360 km until its confluence with the Paranaíba River, where both coalesce onto the Paraná River (a major component of the La Plata Basin). It is an illustrative example of thoroughly impounded river, with 12 large dams, with the Camargos hydroelectric power plant being the last dam upbound, implemented around 1960. It is closely followed downstream by another dam, from the Itutinga power plant, operational since 1955, which delimits a singular, relatively small closed impoundment (without tributaries). There, an environmental hatchery used to actively operate for decades in broodstock production and release, mainly of *P. lineatus*, throughout the region, until 2018. It provides a singularly convenient stage for testing for the genetic effects of broodstoking. Next, further downstream, lies the relatively new Funil power plant dam (constructed circa the year 2000), the only of these dams accommodating a fish transposition mechanism, a lift (Suzuki *et al*., 2011). These three dams are featured in Figure 1. The degree in which these large dams genetically isolate *P. lineatus* in the Upper Grande River is not currently known, nor are the genetic effects from broodstock release and fish transposition operations there.

**Figure 1.**
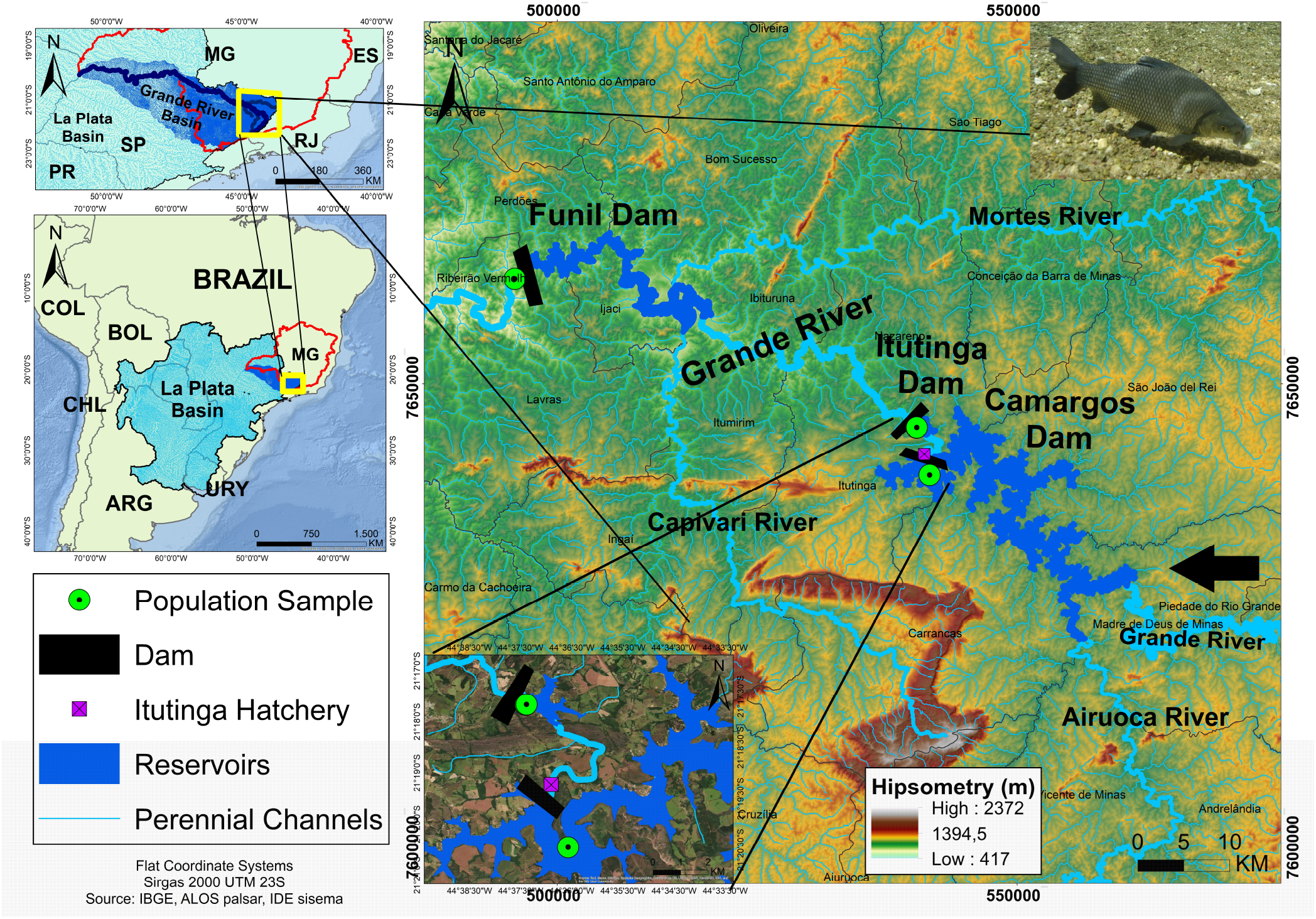
Geographical location of the studied population samples for curimba *Prochilodus lineatus* from the Upper Grande River Basin, MG, Brazil. Photography by André Seale.

Only a handful of genomic resources are currently available for prochilodontid fish species. The mitochondrial DNA sequence for *P. lineatus* was unveiled by Carmo *et al*. (2016), while other *Prochilodus* species’ mitogenomes were presented by Chagas *et al*. (2016) and Santos *et al*. (2021). Regarding the nuclear genome, Stornioli *et al*. (2021) have disclosed NGS data for *P. lineatus* and Yepes-Blandón *et al*. (2022), in turn, have delivered the first draft of a full reference genome for *Prochilodus magdalenae*, a Colombian species, although not publicly releasing NGS data.

Our core objectives in this work were *i)* testing the hypothesis of the occurrence of mixed stocks across *P. lineatus* populations segregated by the last three upbound dams from the Upper Grande River, MG, Brazil; *ii)* testing the hypothesis that dams are genetically isolating migratory fish in the region and; *iii)* testing the hypothesis that broodstocking practices have altered genetic diversity in populations of fish trapped between dams. We aimed at doing so through the development of a genome-wide set of microsatellite *loci*, for rapid marker development in *P. lineatus*, while making available new genomic resources for this fish, a model for Neotropical freshwater migratory species in general.

## Material and Methods

### Ethics statement

All proceedings presented here were conducted under the guidance and licenses 27/2011 and 6991130223 of the Committee on Ethics in Animal Use (CEUA-UFSJ); along CGEN licences A9D0E5 and AC5A784; and SISBIO licenses 37222-2 and 85497-1. No experimental assays were conducted on live specimens.

### Genomics, bioinformatics and microsatellite markers development

Genomic libraries of whole genome shotgun fragments (500 bp) were produced for NGS, out of a single adult female individual from the Itutinga Hatchery Station (21°19’20.7”S 44°36’56.9”W - Figure 1. The methodological steps and parameters for NGS and initial microsatellite characterization were the same as presented in Yazbeck *et al*. (2018), and carried out by an external service provider (SP), BGI (Tai Po, Hong Kong). The resulting 90 bases paired-end (PE) short-reads were organized as three sets of twin FASTQ files. We verified the quality of DNA short-reads using MultiQC (Ewels *et al*., 2016), and computed all of its 21-mers (k-mer substrings) in Jellyfish (Marçais and Kingsford, 2011). These latter results were applied in the estimation of the expected genome size of *P. lineatus* through a frequency histogram, in R version 4.4.1 (R Core Team, 2024).

We used all the DNA short-reads, along a set of fasta format contigs containing microsatellites resolved by the SP upon the original undisclosed assembly, to execute a *de novo* genomic assembly, ourselves. For this step, we used MEGAHIT (Li *et al*., 2015), using k-mer=47, to generate new contigs and, then, submitting these to the SOAPdenovo-fusion module from SOAPdenovo 2 (Luo *et al*., 2012), to resolve scaffolds. This genome assembly was processed in the Sagarana high-performance computing cluster (CEPAD-ICB-UFMG, Belo Horizonte, Brazil). The National Center for Biotechnology Information’s (Sayers *et al*., 2022) contamination screening tool FCS (Astashyn *et al*., 2024) was used by NCBI staff and ourselves for filtering out mitochondrial assembly, sequences from microorganisms and library adaptors found in the final scaffold level sequence. Assembly statistics were accessed through assemblyStats (http://gif.biotech.iastate.edu) and its visualization by assembly-stats 17.02 (Challis, 2017).

We mapped the microsatellites characterized in the steps above in the new genomic assembly with the strategy also delineated in Yazbeck *et al*. (2018). A sequence alignment map (SAM) was then produced with BBmap (Bushnell, 2014), by mapping short-reads unto the genomic assembly, and finally converted to a binary format (BAM) with SAMtools (Li *et al*., 2009).

For this work we developed new species-specific microsatellite markers to be applied in the population genetics analysis. We used the results of the genomic microsatellite *loci* search by the SP for choosing candidate *loci* for empirical tests. The microsatellites were sorted by decreasing motif size and number of repeats, in order to maximize chances of finding polymorphic *loci* (Brandström and Ellegren, 2008). A total of 22 candidates primer pairs were picked (12 from the top sorted candidates and 10 at random) and then synthesized by an SP (Integration DNA Technologies, Coralville, USA). These were applied to wet bench assays in four individual DNA samples. Polymerase chain reaction (PCR) was carried in a final volume of 10 µl, with initial conditions of 1 µl of unquantified chelex extracted genomic DNA to seed the reaction; 10 pmol of each primer; MgCl_2_ (2 mM); Tris-HCl (10 mM); KCl (50 mM); and 1 unity of Taq DNA polymerase enzyme (Phoneutria, Belo Horizonte, Brazil). Thermal PCR cycles were as following: initial genomic DNA denaturation at 94ºC for 4 min.; 30 iterated cycles of denaturation at 94ºC for 30 s, primers annealing temperature (specific for each primer pair) for 30 s, and amplicon elongation step at 72ºC for 30 s; followed by a final elongation step, at the same temperature, for 5 min.

The resulting PCR products were visualized by polyacrylamide gel electrophoresis at 10%, at 15 V/cm, loaded along with 25 bp DNA fragment size ladder, stained with ethidium bromide. *Loci* yielding polymorphic patterns were then assayed in a 20 individuals sample, for optimization of PCR conditions. Gels were photographed under UVB light, and scored with GelAnalyzer. Scored DNA fragments were then converted to nominal allele classes, through a in house semi-automated (*i*.*e*. automated, under human supervision) binning approach, based on frequency histograms and formatted for downstream analysis.

### Population genetics

A total of 100 *P. lineatus* individual tissue samples (ethanol preserved fin fragments) were used, from the DNA bank of the Laboratório de Recursos Genéticos (LARGE-UFSJ). These were collected in 2013 from three locations (Figure 1) from the Upper the Grande River, MG: downstream the Funil dam, upstream the Itutinga dam and upstream the Camargos dam (see Suzuki *et al*. 2011 for a more detailed characterization from these locations). An additional sample of 40 individuals was obtained from the Itutinga dam reservoir in 2023 (Table 1).

**Table 1.** Evaluated *Prochilodus lineatus* individuals sampled, from the last three dams from the Upper Grande River, MG, Brazil.

| Sample | ID | Location | Year | N |
| --- | --- | --- | --- | --- |
| **Camargos** | 1-20 | 21°20'27.1"S 44°36'34.3"W | 2013 | 20 |
| **Itutinga 1** | 21-60 | 21°17'39.9"S 44°37'24.1"W | 2013 | 40 |
| **Itutinga 2** | 61-100 | 21°17'30.4"S 44°37'11.4"W | 2023 | 40 |
| **Funil** | 101-140 | 21°08'39.4"S 45°02'19.1"W | 2013 | 40 |
| **Total** |  |  |  | **140** |

ID=Individual label; N=Sample size.

Following DNA marker development experiments, a total of 15 new empirically validated microsatellite *loci* were applied to 140 individuals of *P. lineatus*. Hardy-Weinberg equilibrium (HWE) and linkage disequilibrium (LD) were tested through the methods implemented in GenePop 007 (Rousset, 2008), for each population sample, with default options for Markov chain Monte Carlo (MCMC) parameters. These were analysed under new significance threshold values for multiple tests, following sequential Bonferroni corrections. Microchecker (van Oosterhout *et al*., 2004) was used for controlling scoring errors (stuttering, large allele dropout and null alleles). Genetic data handling, conversions and table visualizations were mainly conducted with the adegenet package for R (Jombart, 2008).

Other genetic diversity and structure estimates for the population samples were obtained through GenAlex (Peakall and Smouse, 2012), including expected and observed heterozygosities (H_E_ and H_O_, respectively); allelic richness, *i*.*e*. number of alleles (N_A_); Jost’s D; the fixation index G”_ST_; and analysis of molecular variance (AMOVA), each with 999 random permutations for assessing probability values. For D and G”_ST_, standard errors (SE) were calculated by jackknifing over *loci* and 95% confidence intervals (CI) were obtained by bootstraping over *loci* (Peakall and Smouse, 2012). Resampling jackknife estimates of total allelic richness, per population samples, were conducted with the aid of EstimateS (Colwell and Elsensohn, 2014) and its confidence intervals were calculated in

R. Rarefaction analysis of allelic richness was conducted for controlling sample size bias, with the aid of R packages dplyr, tidyr and pegas (Paradis, 2010). R visualizations were plotted with the aid of ggplot2.

A discriminant analysis of principal components (DAPC - Jombart *et al*., 2010) was conducted through adegenet, by *de novo* grouping, initially retaining 80% of total variation captured by principal components (PC) and with optimal k chosen according to the Bayesian information criterion (BIC). A cross-validation procedure, with 100 replicates, was executed and used to gauge the DAPC analysis reliability, with the best number of PC found. Finally, the DAPC was also conducted with population sample information, with the same optimal number of PC retained in cross-validation, in order to visually record the genetic diversity within and among groups.

### Hierarchical analysis of the genetic population structure

The genotypic data was also processed with analytical tools implemented in PopCluster Wang (2022). A preliminary maximum likelihood estimation (MLE) analysis is performed upon a null-model for admixture (*i*.*e*. a strict mixture model, where an individual composite genotype is assigned exclusively to a single ancestral deme), for the clustering individuals into K demes. It takes advantage of an annealing algorithm, which prevents estimates from being trapped within a sub-optimal value peak, enhancing its chances of visiting of all peaks in the parameter space and, thus, increasing the likelihood of finding its global maximum regarding the data. These results are then used as reference for performing MLE of individual admixture coefficients. This allows for the refinement of the assignment of individuals to one of the K possible population clusters, without information (supervision) from the sample’s origin. Unequal allele frequencies were assumed and the weak scaling factor was used. The choice of the optimal K value explaining sample substructure (K=1-14, N=100 replicate runs) was primarily based on the 2^nd^ order rate of log-likelihood change estimator, D_LK2_, along computations of the F_STIS_ estimator (Wang, 2022). This initial procedure was labelled as a 1^st^ order analysis of population genetic structure. Then, to explore possible hierarchically hidden strata of genetic structure (Vähä *et al*., 2007; Janes *et al*., 2017), the best replicate run results for the chosen K value were used for rearranging samples into new clusters. These were each, in turn, independently subjected to a 2^nd^ order admixture analysis, ranging from K=1-7. This was reiterated once more, as a 3^rd^ order analysis of population structure, assaying K=1-4. The resulting rearranged clusters, across hierarchical orders, were examined for genetic diversity and structure measurements, as the original population samples.

We also performed a parallel Bayesian inference, using the STRUCTURE suite (Pritchard *et al*., 2000), for the admixture model, *de novo*, with different priors for each population. A total of 5 × 10^5^ steps were discarded before computing 1 × 10^6^ MCMC iterations, with K varying from 1 to 8 and 20 replicates each. Estimation of best K value for the STRUCTURE analysis was carried primarily with the estimators implemented in Kfinder (Wang, 2019) and visualization was conducted with the help of clumpak (Kopelman *et al*., 2015).

## Results

### Genomics, bioinformatics and microsatellite markers development

Approximately 483.7 million short-reads (43.53 gigabases) were generated for *P. lineatus* and deposited at the the Sequence Read Archive (SRA) database, from NCBI (Bioproject accession code PRJNA1120875). Its CG content was 42% and per base quality value (Q) ranged from 31 to 39, with an average of 38 per read. Based on the k-mer counts, total genome size was estimated at 1.65 Gb.

The *de novo* genome assembly generated was 1.4 Gb, with 31X coverage. This Whole Genome Shotgun project has been deposited at DDBJ/ENA/GenBank under the accession JBGFTF000000000. The version described in this paper is version JBGFTF010000000. The total number of contigs/scaffolds was 738,832 sequences, with N_50_=5,503 bp and L_50_=52,608. The average length of contigs was 1,901 with the longest being 192,837 bp (Figure 2). The BAM file associated with it can also be retrieved from NCBI (PRJNA1120875). A total of 37,476 perfect di-to hexa-nucleotide microsatellite *loci* for *P. lineatus* were characterized *in silco* and compiled in Supporting Information (Supplementary Files 1 and 2). These were mapped upon the genome assembly, but around 12% of putative *loci* could not be traced back to it.

**Figure 2.**
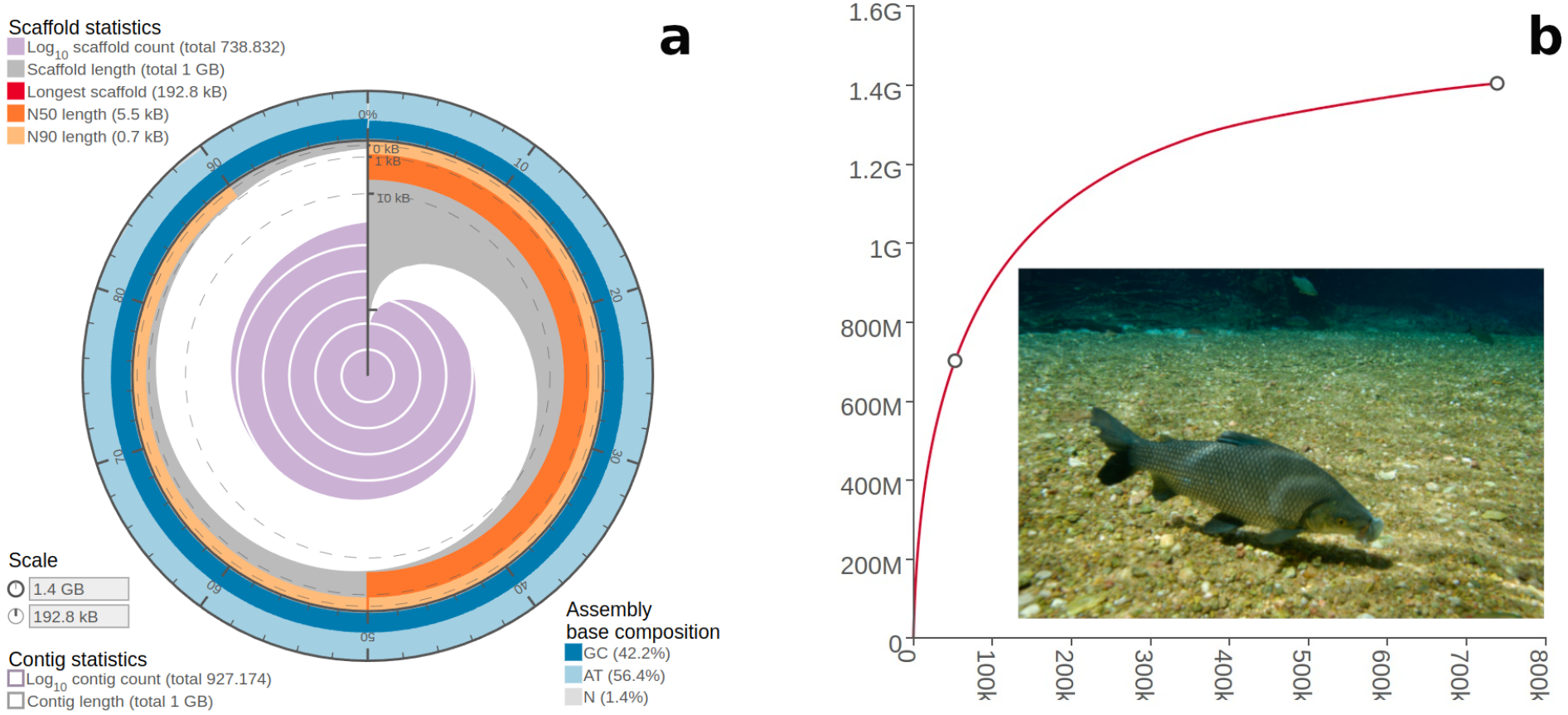
Genome assembly statistical features for *Prochilodus lineatus*. **a**) The radius inside the circular plot marks the length of the longest scaffold resolved and the grey segments represent the cumulative percentage of the assembly contained in scaffolds equal or longer than a particular length. **b**) Cumulative contig/scaffold length of the genomic assembly, as a function of the the number of sequences resolved. The point in the middle of the curve marks the L_50_ feature, the smallest number of sequences, which the sum surpass half the total genome assembly length (point at the end of the cuve). G=Gigabases and k=thousand of sequences.

Of the 22 primer pairs tested, five (Prol14, Prol15, Prol18, Prol31 and Prol52) resulted in complex amplification patterns and two (Prol56 and Prol28) failed to amplify. Fifteen new microsatellite markers were successfully amplified and results for the pooled sample of 140 fish are presented in Table 2. The optimal PCR conditions were as initially tested, except for Prol02 and Prol19 (6.7 pmol of each primer), the latter using 35 cycles, and Prol54, which also used 35 cycles. The average number of alleles per locus was 22.7 (±1.3 standard error - SE).

**Table 2.**
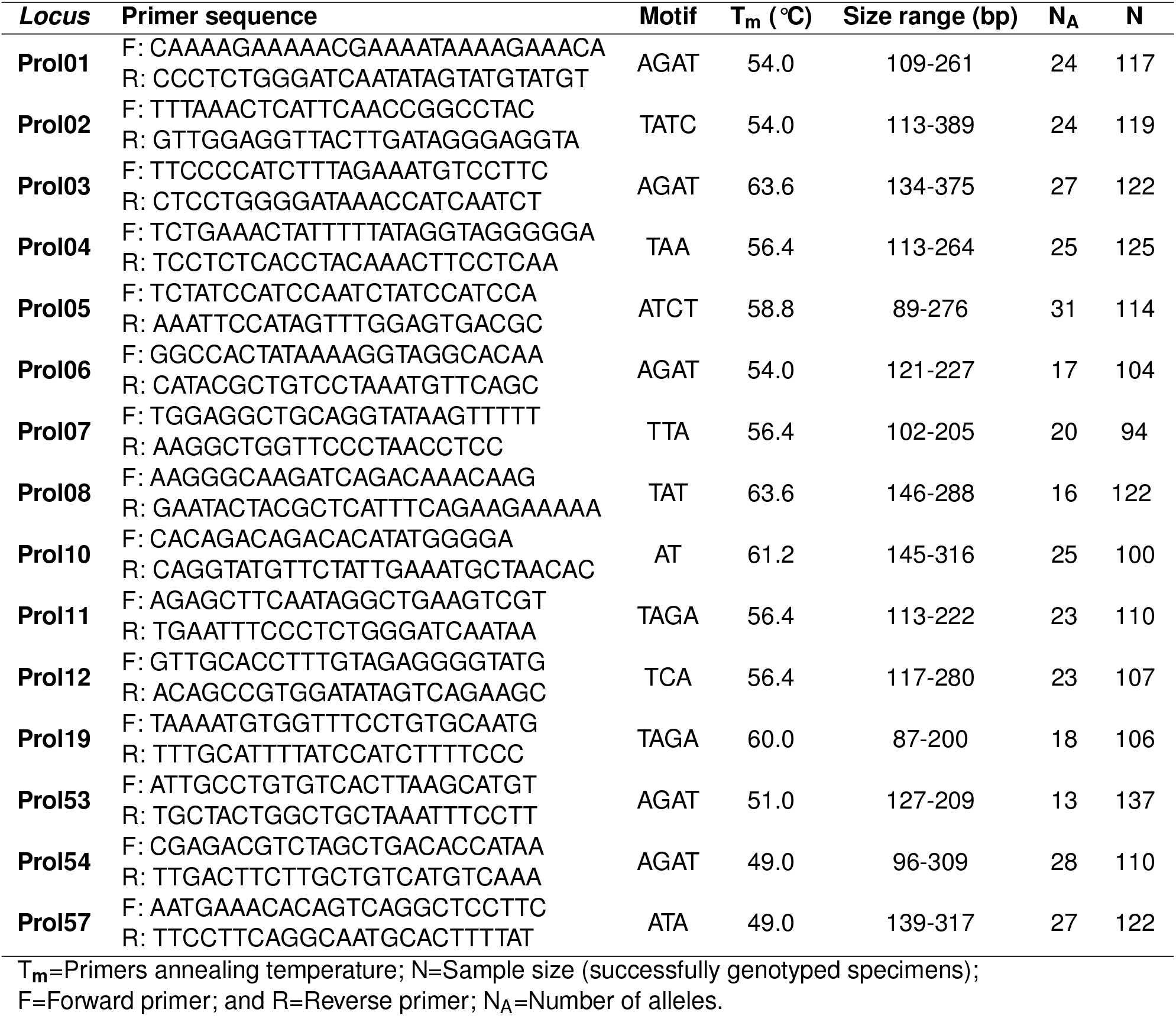
New validated microsatellite markers for *Prochilodus lineatus*, along observed amplified DNA fragment length intervals for each locus and its respective defined allelic classes.

The amplification/genotyping failure rate varied from 2.1% (Prol53) to 32.8% (Prol07), with average 18.6% (±7.9 SE), including instances where one allele was scored and the other deemed undetermined for the same diploid genotype. Six DNA samples (four from Camargos, one from Itutinga 1 and one from Funil) failed to amplify between five and ten *loci*. These were removed from data set and most downstream analyses were performed with and without it, in parallel. Marginal or no differences were observed when comparing results and, thus, these samples were retained.

### Population genetics

Adjusted significance for 60 multiple HWE tests was set to p<0.008 following the sequential Bonferroni correction. By this criterion, only six *loci* were deemed in HWE. The only locus found to conform with HWE within every sample, Prol07, was not in equilibrium across all four population samples (p=0.002). All deviations were towards heterozygote deficits. Tests for LD within population samples, had an adjusted significance level *α*=0.0001. No LD was inferred, except for six instances: Prol06 and Prol54 for Camargos; Prol01 and Prol19, Prol05 and Prol57 for Itutinga 1; Prol01 and Prol06, Prol04 and Prol07 for Itutinga 2; and finally, Prol53 and Prol57 for Funil. The following *loci* pairs exhibited LD for the pooled sample of fish: Prol01 and Prol06; Prol04 and Prol07; Prol01 and Prol19; Prol06 and Prol54; Prol05 and Prol57; and lastly, Prol53 and Prol57. There was no evidence for scoring errors due to stuttering or large allele dropout, but the presence of null alleles was suggested as possible, due to general excess of homozygotes for most allele size classes, for all *loci*, in all population samples, except for Prol07.

Allelic richness ranged from 10.8 (±0.8 SE) alleles for Camargos to 16.9 (±0.9 SE) alleles for and Itutinga 2 (Table 3). This trend was confirmed by the jackknife estimates of allelic richness (Supporting Information - Supplementary Figure 1), along the results for the rarefaction analysis (Supplementary Figure 2). Camargos also had the lowest expected heterozygosity, while displaying the highest inbreeding coefficient value, along the Itutinga 2 sample (Table 3). All population samples exhibited private alleles, ranging from 7 alleles for Camargos and 37 alleles for Itutinga 2 (average 19.7 ±12.7 SE).

**Table 3.** Genetic diversity results for each microsatellite maker across population samples in *Prochilodus lineatus*. Probability for Hardy-Weinberg test had significance set to *α*=0.008. All probability values (p) are significant, except where noted (ns), and values deemed zero are actually extremely low non-zero values p<0.00009.

| Sample | Locus | N | N<sub>A</sub> | H<sub>O</sub> | H<sub>E</sub> | F | HWE (p) |
| --- | --- | --- | --- | --- | --- | --- | --- |
| Camargos | Prol01 | 17 | 14 | 0.65 | 0.91 | 0.29 | 0.0023 |
|  | Prol02 | 15 | 14 | 0.60 | 0.89 | 0.33 | 0 |
|  | Prol03 | 13 | 13 | 0.31 | 0.91 | 0.66 | 0 |
|  | Prol04 | 17 | 10 | 0.41 | 0.87 | 0.53 | 0 |
|  | Prol05 | 14 | 13 | 0.36 | 0.9 | 0.6 | 0 |
|  | Prol06 | 14 | 9 | 0.43 | 0.81 | 0.47 | 0 |
|  | Prol07 | 15 | 10 | 0.73 | 0.85 | 0.14 | 0.0191<sup>ns</sup> |
|  | Prol08 | 13 | 4 | 0.08 | 0.65 | 0.88 | 0 |
|  | Prol10 | 16 | 10 | 0.69 | 0.85 | 0.19 | 0.0049 |
|  | Prol11 | 17 | 12 | 0.35 | 0.88 | 0.6 | 0 |
|  | Prol12 | 13 | 6 | 0 | 0.77 | 1 | 0 |
|  | Prol19 | 12 | 9 | 0.50 | 0.84 | 0.41 | 0.0004 |
|  | Prol53 | 18 | 9 | 0.33 | 0.77 | 0.57 | 0 |
|  | Prol54 | 17 | 15 | 0.41 | 0.91 | 0.55 | 0 |
|  | Prol57 | 17 | 11 | 0.41 | 0.87 | 0.53 | 0 |
|  | Over *loci* | 15.2 | 10.6 | 0.42 | 0.85 | 0.52 | 0 |
| Itutinga 1 | Prol01 | 33 | 16 | 0.27 | 0.91 | 0.7 | 0 |
|  | Prol02 | 33 | 18 | 0.55 | 0.92 | 0.41 | 0 |
|  | Prol03 | 31 | 16 | 0.52 | 0.91 | 0.43 | 0 |
|  | Prol04 | 36 | 13 | 0.31 | 0.9 | 0.66 | 0 |
|  | Prol05 | 26 | 22 | 0.62 | 0.94 | 0.34 | 0 |
|  | Prol06 | 34 | 11 | 0.21 | 0.87 | 0.76 | 0 |
|  | Prol07 | 20 | 12 | 0.95 | 0.88 | -0.08 | 0.0157<sup>ns</sup> |
|  | Prol08 | 38 | 10 | 0.45 | 0.85 | 0.48 | 0 |
|  | Prol10 | 33 | 20 | 0.73 | 0.94 | 0.22 | 0 |
|  | Prol11 | 30 | 16 | 0.57 | 0.89 | 0.37 | 0 |
|  | Prol12 | 31 | 13 | 0.16 | 0.9 | 0.82 | 0 |
|  | Prol19 | 30 | 13 | 0.23 | 0.91 | 0.74 | 0 |
|  | Prol53 | 39 | 9 | 0.31 | 0.64 | 0.52 | 0 |
|  | Prol54 | 34 | 20 | 0.38 | 0.92 | 0.59 | 0 |
|  | Prol57 | 38 | 16 | 0.47 | 0.91 | 0.48 | 0 |
|  | Over *loci* | 32.4 | 15 | 0.45 | 0.89 | 0.50 | 0 |
| Itutinga 2 | Prol01 | 34 | 19 | 0.29 | 0.92 | 0.68 | 0 |
|  | Prol02 | 36 | 18 | 0.56 | 0.91 | 0.39 | 0 |
|  | **Prol03** | 39 | 19 | 0.36 | 0.93 | 0.61 | 0 |
|  | **Prol04** | 33 | 14 | 0.27 | 0.91 | 0.7 | 0 |
|  | **Prol05** | 38 | 24 | 0.39 | 0.94 | 0.58 | 0 |
|  | **Prol06** | 27 | 14 | 0.11 | 0.89 | 0.87 | 0 |
|  | **Prol07** | 27 | 18 | 0.81 | 0.91 | 0.1 | 0.0498<sup>ns</sup> |
|  | **Prol08** | 39 | 14 | 0.62 | 0.87 | 0.3 | 0 |
|  | **Prol10** | 27 | 15 | 1 | 0.91 | -0.1 | 0 |
|  | **Prol11** | 33 | 17 | 0.27 | 0.88 | 0.69 | 0 |
|  | **Prol12** | 32 | 17 | 0.34 | 0.91 | 0.62 | 0 |
|  | **Prol19** | 31 | 13 | 0.13 | 0.89 | 0.86 | 0 |
|  | **Prol53** | 40 | 10 | 0.18 | 0.65 | 0.73 | 0 |
|  | **Prol54** | 34 | 19 | 0.29 | 0.92 | 0.68 | 0 |
|  | **Prol57** | 36 | 22 | 0.81 | 0.92 | 0.13 | 0.0210<sup>ns</sup> |
|  | **Over loci** | 33.7 | 16.9 | 0.43 | 0.89 | 0.52 | 0 |
| **Funil** | **Prol01** | 33 | 16 | 0.15 | 0.92 | 0.84 | 0 |
|  | **Prol02** | 35 | 17 | 0.49 | 0.91 | 0.46 | 0 |
|  | **Prol03** | 39 | 21 | 0.51 | 0.95 | 0.46 | 0 |
|  | **Prol04** | 39 | 17 | 0.51 | 0.92 | 0.44 | 0 |
|  | **Prol05** | 36 | 27 | 0.56 | 0.95 | 0.42 | 0 |
|  | **Prol06** | 29 | 12 | 0.21 | 0.79 | 0.74 | 0 |
|  | **Prol07** | 32 | 11 | 0.91 | 0.85 | -0.07 | 0.4959<sup>ns</sup> |
|  | **Prol08** | 32 | 13 | 0.66 | 0.86 | 0.24 | 0 |
|  | **Prol10** | 24 | 18 | 1 | 0.93 | -0.08 | 0.7323<sup>ns</sup> |
|  | **Prol11** | 30 | 18 | 0.4 | 0.92 | 0.57 | 0 |
|  | **Prol12** | 31 | 19 | 0.29 | 0.9 | 0.68 | 0 |
|  | **Prol19** | 33 | 15 | 0.15 | 0.91 | 0.83 | 0 |
|  | **Prol53** | 40 | 11 | 0.23 | 0.68 | 0.67 | 0 |
|  | **Prol54** | 25 | 14 | 0.72 | 0.91 | 0.2 | 0.0052 |
|  | **Prol57** | 31 | 21 | 0.71 | 0.93 | 0.23 | 0 |
|  | **Over loci** | 32.6 | 16.67 | 0.50 | 0.89 | 0.44 | 0 |
|  | **Overall** | 140 | 341 | 0.45 | 0.91 | 0.50 | 0 |

N=Sample size; N<sub>A</sub>=Number of alleles; H<sub>O</sub>=Observed heterozygosity; H<sub>E</sub>=Expected heterozygosity; F=Inbreeding coefficient; HWE=Exact test for Hardy-Weinberg equilibrium.

The AMOVA results showed that only 1% of total variation was explained by the four population samples, with the remaining variation being explained by differences among individuals (52%) and within genomes (47%). All pairwise fixation estimates for G”_ST_ and Jost’s D were significantly different from 0, including between samples collected in the same location, but at different times (Table 4). These Itutinga samples (2013 and 2023) had the lowest values, while Camargos showed the highest overall pairwise variation and differentiation, for both respective indices. Considering all 15 microsatellite *loci*, among the four population samples, the unbiased standardise fixation index G”_ST_ reached 0.16 (±0.03 SE; 0 to 0.26 CI, n=999; p=0.001) and the total genetic differentiation estimate of Jost’s D, was 0.15 (±0.03; 0 to 0.24 CI, n=999; p=0.001).

**Table 4.** Summary of pairwise fixation index G”ST (below diagonal) and Jost’s genetic distance D (above diagonal) for microsatellites diversity among population samples of *Prochilodus lineatus* from the Upper Grande River, MG.

|  | Camargos | Itutinga 1 | Itutinga 2 | Funil |
| --- | --- | --- | --- | --- |
| **Camargos** | - | 0.156** | 0.222** | 0.203** |
| **Itutinga 1** | 0.171** | - | 0.066* | 0.087** |
| **Itutinga 2** | 0.241** | 0.072* | - | 0.167** |
| **Funil** | 0.221** | 0.095** | 0.180* | - |

### Hierarchical analysis of the genetic population structure

Maximum likelihood estimation (MLE) of admixture (Figure 3) pointed to K=2 as the most likely number of clusters, according to Wang’s D_LK2_ metric. However, F_STIS_ suggested K=4 as the most likely number of groups (Supporting Information - Supplementary Figure 3). The 1^st^ order admixture analysis allowed the definition of two clusters (with respective individual IDs - Figure 3a). Estimates of D_LK2_ for both these clusters yielded the value of K=2 as optimal in the 2^nd^ order analysis (Figure 3b), while F_STIS_ recorded values of K=5 and K=7 (Supplementary Figure 4). Finally, these four newly defined individual rearrangements were studied in the 3^rd^ order analysis (Figure 3c), and D_LK2_ consistently pointed to K=2 for each and every subject cluster (Supplementary Figure 5; individual assignments for all hierarchical orders can be found in Supplementary File 3). We addressed and discarded the possibility of panmixia (K=1), at this and each hierarchical order level, by directly inspecting log likelihood values of the clustering analysis, against the results for the assumed value for K (Supporting Information - Supplementary File 4 - Supplementary Table 1).

**Figure 3.**
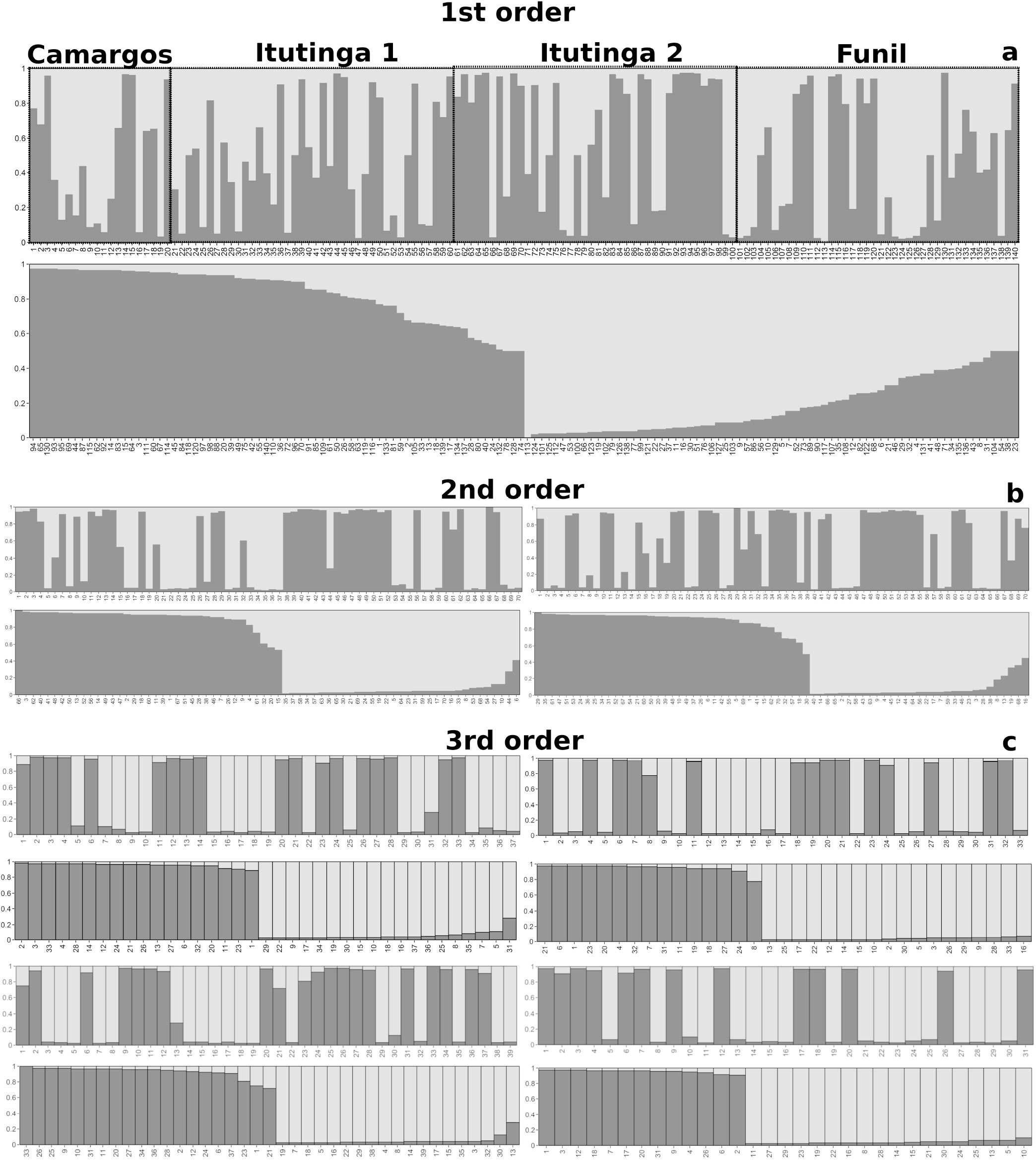
Individual admixture coefficients (bars) for K=2, across inclusive levels of hierarchical genetic structure of *Prochilodus lineatus* populations from the Upper Grande River, MG, Brazil. Different shades of grey represent alternative clusters of origin, with the proportional size of bars indicating the respective genomic contributions from ancestral demes. For each panel, the diagram above shows the sample in the original input sequence, while diagram below shows the sample organized as newly defined clusters. **a**) 1^st^ order analysis, showing fish grouped according to population sample and original ID. **b**) 2^nd^ order analysis from clusters defined in the 1^st^ order. **c**) 3^rd^ order analysis of 2^nd^ order clusters. For 2^nd^ and 3^rd^ orders, the numbers below bars are arbitrary and not equivalent to original ID.

These different MLE sample arrangements showed a consistent decrease in intrapopulation diversity parameters for the higher orders, such as expected heterozygosity and number of alleles (Supplementary File 4 - Supplementary Table 2), while interpopulation indicators (G”_ST_ and Jost’s D) steadily increased (Supplementary File 4 - Supplementary Tables 3-5). The HWE analysis of MLE defined clusters indeed showed a tendency towards Hardy-Weinberg expectations in higher order group arrangements. In average, 3^rd^ order clusters had more *loci* in HWE, with one particular cluster showing eight *loci* in equilibrium (Supplementary File 5).

The DAPC performed without population sample information pointed for two main clusters, according to the BIC (Figure 4, Supplementary Figure 6). This initial DAPC also revealed that two alleles from Prol53 (03 and 04) had the most weight in defining variable contributions. The cross-validation assay pointed to 30 as the ideal number of PC to be retained (Supplementary Figure 7) and the ratification of the DAPC according to this parameter led to very convergent results (Supplementary Figure 8). The DAPC performed with this number of PC and information of population samples revealed an apparent level of genetic differentiation among locations (Figure 5).

**Figure 4.**
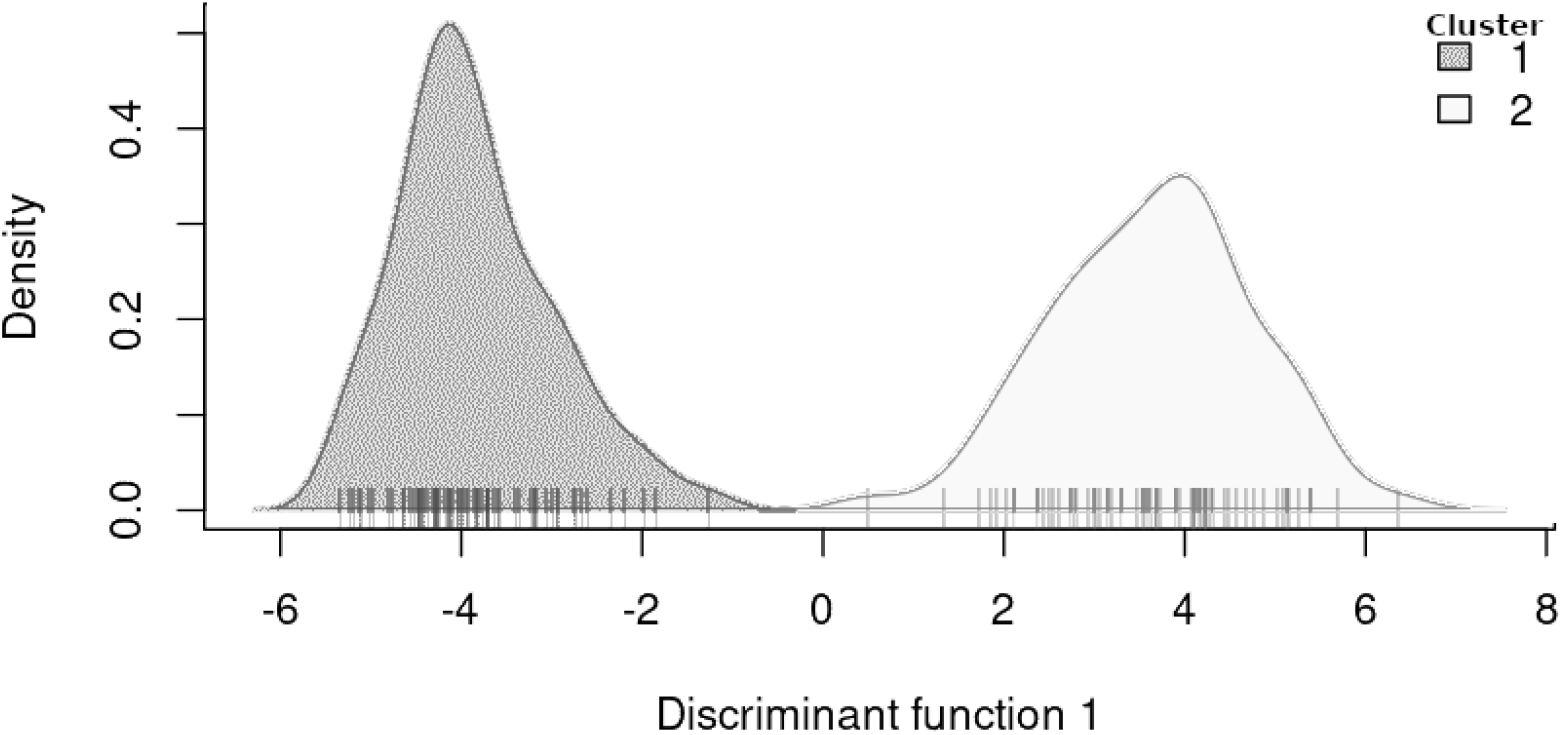
Density of individual clusters defined by the initial *de novo* discriminant analysis of principal components (DAPC) for *Prochilodus lineatus* from the Upper Grande River, MG.

**Figure 5.**
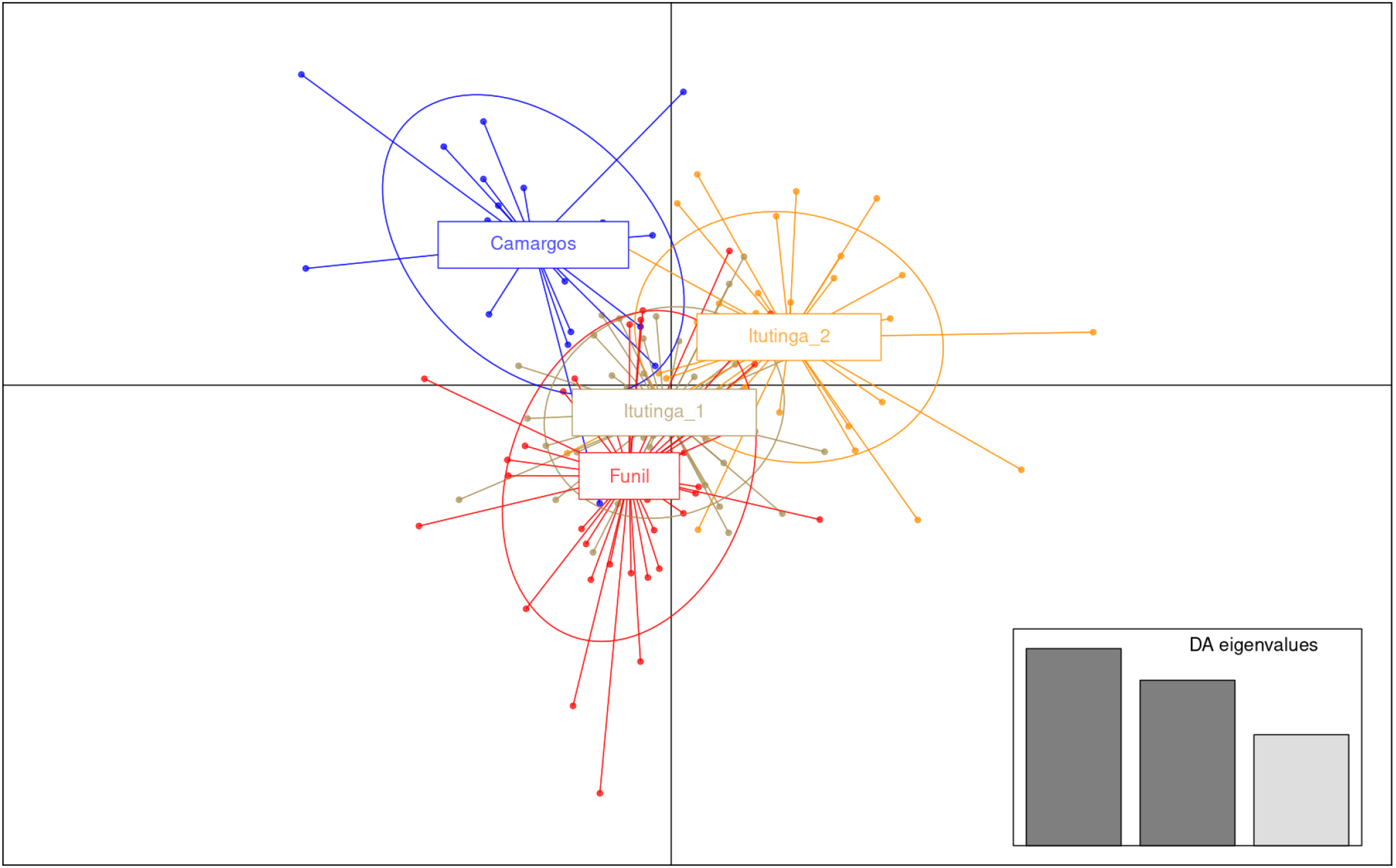
DAPC for *Prochilodus lineatus* from the Upper Grande River, MG, performed with population sample information, with three discriminant functions (scaled discriminant analysis eigenvalues showed on the bottom right).

The Bayesian approach to admixture also resulted in alternative values for best K estimates, with K=8 according to Pritchard’s Pr(X|K) while, both, Evanno’s ΔK and Wang’s parsimony estimators resolved K=5 (Figure 6).

**Figure 6.**
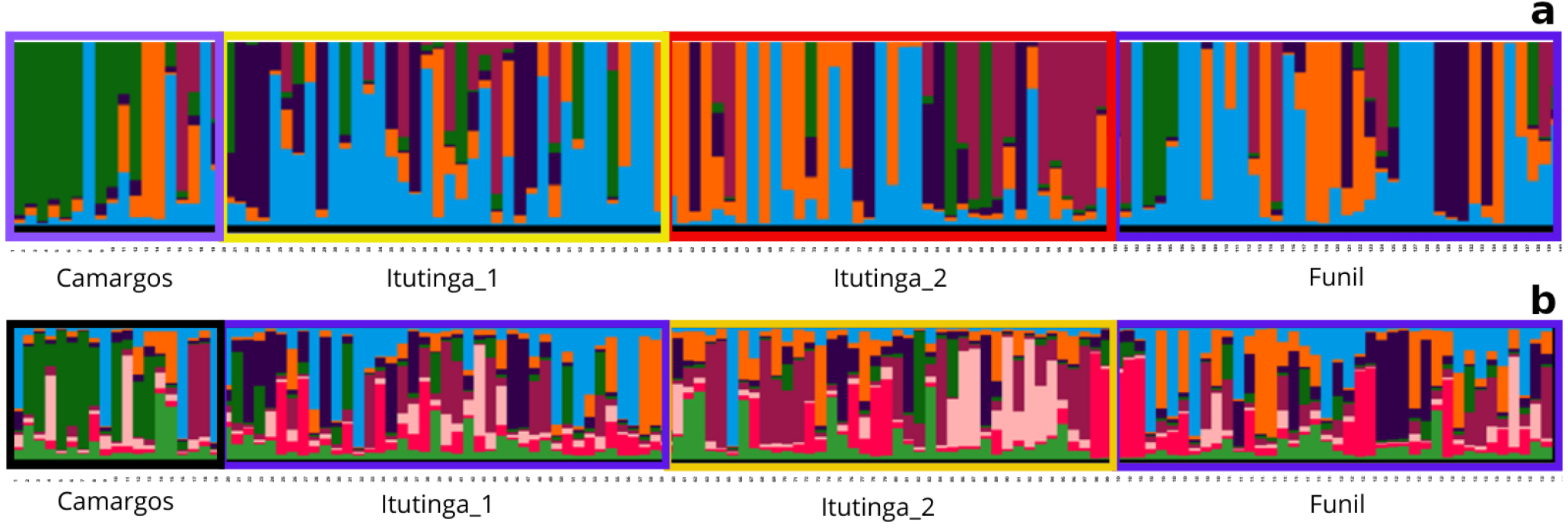
Admixture plots from the STRUCTURE clustering results for a) K=5 and b) K=8 for *Prochilodus lineatus* from the Upper Grande River, MG. Each colour represents a different cluster. Note the apparent differentiation at Camargos and Itutinga 2.

## Discussion

### Genomics, bioinformatics and microsatellite markers development

Genome assemblies are becoming currently mandatory for many different biological disciplines (Jung *et al*., 2020). Our work takes a new step towards this goal, by publicly presenting the first comprehensive genomic resources for this highly iconic species, *P. lineatus*. The volume of short-read DNA sequence data presented here surpass the only previously available FASTQ data for *P. lineatus* (Stornioli *et al*., 2021) by a factor of 12. This number of sequences is, though, one order of magnitude below the short-reads data reported for the trans-Andean congeneric species, *P. magdalenae* (Yepes-Blandón *et al*., 2022) which, however, is not currently publicly available. The CG content and quality rate of reads were the same as the ones previously presented for other piracema fish (*e*.*g*. Yazbeck *et al*., 2018; Graciano *et al*., 2022). It’s also the most extensive dataset of short-reads publicly disclosed so far for any member of the Prochilodontidae family.

This work provides the first *de novo* genome assembly for *P. lineatus*. It’s slightly longer then the described genome size reported in *P. magdalenae* (Yepes-Blandón *et al*., 2022), despite its extensive fragmentation, and shorter than estimated by the k-mer method carried, which is not always precise (Pflug *et al*., 2020). The limited length of the short-reads used here contributed to curtailing our genome assembly, primarily motivated by the exclusive search of microsatellite *loci*. The full sequence of this *P. lineatus* specimen’s mitogenome was previously presented in Santos *et al*. (2021). This novel genome assembly might be explored for the initial annotation of longer characterized sequences (*e*.*g*. Graciano *et al*., 2021).

The genome wide microsatellite panel produced here, with more than 37 thousand putative *loci*, can now be free and easily accessed for the rapid, low-cost development of potentially hundreds of new empirically validated markers. It may also be explored for technological development of practical tools for hatcheries and management of impacted areas, such as multiplexed sets of markers (*e*.*g*. Carvalho *et al*., 2019). Some of these resources may also, arguably, be applied in different fishes, particularly in other prochilodontid and characid species (*e*.*g*. Barbosa *et al*., 2006; Carmo *et al*., 2015). The relatively high rate of success in empirical assays attests to the high quality of this panel. These 15 new DNA markers presented here increase by 84% the number of available microsatellites for *P. lineatus*.

We observed a variable rate of amplification success among these new markers. Some *loci* presented a relatively high amplification/genotyping failure rate (*e*.*g*. Prol07). Some approaches (*e*.*g*. admixture assignments) are found to be reasonably insensible to missing data, specially when it is not due to a systematic cause (Falush *et al*., 2007; Wang, 2022). Even null alleles might have only a marginal impact (Carlsson, 2008; Putman and Carbone, 2014; Budowle and Sajantila, 2024). For some other applications, the issue of missing data is more critical (*e*.*g*. estimating effective population size, Ne – Peel *et al*., 2013). The microsatellite Prol53 had a particularly high rate of amplification and genotyping and high information content. Despite Microchecker suggesting the possible presence of null alleles for most *loci* in all populations, we consider this an artefact due to the inherent sample substructuring, which leads to critical confounding effects in this analysis (Waples, 2015).

### Population genetics

A study by Wang *et al*. (2021) suggests that applying less than 11 or 12 microsatellite markers may lead to important bias in population genetics structure assessments. The results presented here are the most extensive efforts so far, concerning the number of analysed *loci*, in *P. lineatus*. It revealed high levels of microsatellite diversity in fish from the uppermost portion of the Grande River, MG, under the influence of large dams. This is a common result to other former studies regarding population genetics in *P. lineatus* with close genomic sampling efforts (*e*.*g*. Rueda *et al*., 2013; Perini *et al*., 2021).

The genetic diversity analysis of population samples showed a lower genetic richness for fish upstream the Camargos dam, the last artificial obstacle towards the upper end of the Grande River Basin. This is an area deemed as priority for conservation of migratory fishes in the state of Minas Gerais (Drummond, 2005) and has been segregated from the downstream system for the last 65 years, since the construction of the Camargos power plant dam. It does not bear a fish passage and larvae and juveniles are not likely to successfully drift downstream (Suzuki *et al*., 2011). Fish transposition, using specimens caught downstream, and repopulation initiatives with hatchery broodstock (mainly *P. lineatus* - Saraiva and Pompeu, 2016) did occur in the region for the last decades, but was mostly executed without purposefully defined goals, other than fulfilling regulatory agencies’ licensing demands or in the context of environmental education actions. This portion of the river is still connected to a last major free-flowing upstream Grande River tributary, the Aiuruoca River, MG. The Camargos dam is, therefore, isolating fish from the rest of the hydrographic system below and stocking operations might have been inefficient in providing appropriate gene flow connectivity for the system above (lack of migrants). Its local populations are most likely undergoing stronger random genetic drift, which leads to genetic impoverishment within demes and differentiation among them. This notion is corroborated by the results showing the largest amount of interpopulational variation and differentiation in Camargos, from all other samples. Thus, our results support the hypothesis that dams are importantly isolating fish populations in the Upper Grande River. The DAPC performed with sample information also supports this notion, based on the visual inspection of microsatellite variation within and among population samples.

The little difference among population samples registered by the AMOVA is compatible with a single, almost panmitic population, or could be explained by more complex, cryptic patterns of genetic variability. Yet, it is deemed as less straightforward in the interpretation of hypervariable markers, such as microsatellites (Jost, 2008). The G”_ST_ fixation index, along the genetic differentiation estimator Jost’s D, pointed to significant deviations from panmixia, with all comparisons significantly different from zero. This supports the hypothesis of isolation by dams, over fish unable to freely migrate.

The 2023 sample from the Itutinga location is the most genetically diverse, followed by fish caught downstream the Funil dam. The latter, as first raised in Yazbeck and Kalapothakis (2007), could be preventing migrating fish of potentially different stocks, from a relatively long downstream stretch of free flowing waters, from proceeding further upstream, despite its fish lift. This virtual *cul-de-sac* could be leading to a higher observed genetic diversity in the sample by accumulation, comparable to the trapped gene pool effect discussed by Perini *et al*. (2021). Upstream this dam lies the mouth from two major free-flowing tributaries of the Grande River, which could be natural candidates for harbouring different *P. lineatus* stocks (*e*.*g*. Suzuki *et al*., 2011), the Mortes River, and the the Capivari River.

The *P. lineatus* population at Itutinga’s reservoir, in turn, has undergone genetic variation and differentiation of around 7% in 10 years. Despite being closed and strongly isolated (without tributaries), it seems to have locally accumulated genetic variation (allelic richness), since 2013. A plausible explanation for this would be the broodstocking operations of *P. lineatus* from the hatchery situated on its banks. Repeated introduction (or escape) of broodstock fish, with varying sources for spawning matrices, along the marked tendency for the occurrence genetic bottlenecks, over particular reproductive efforts (single or few spawned couples), could be yielding alternative sets of new alleles being introduced into this limited stretch of the Grande River. While this location had the highest levels of diversity (N_A_ and H_E_), it also had the highest inbreeding coefficient, reinforcing the notion of hatchery driven substructuring. For more than five years now, though, the hatchery station operates mainly for scientific research and has ceased its fish stocking in the Grande River, MG, and region. These results support the hypothesis that hatcheries can influence the genetic diversity in populations of fish trapped between dams.

Regarding our main hypothesis of the occurrence of mixed stocks in the Upper Grande River, MG, the extensive verified lack of adherence to Hardy-Weinberg proportions in all population samples, always towards heterozygotes deficit, along some verified LD, suggest strong genetic stratification in all sampled populations (*i*.*e*. more than one deme being sampled) and, thus, is in accordance to its prediction. The fact it is so pervasive across *loci* allows us to initially disregard other putative confounding factors, such as eventual null alleles. It can be biologically interpreted as an expression of the Wahlund effect, leading to a pronounced downward departure from the expected heterozygosity under HWE.

### Hierarchical genetic structure analysis

The admixture model seems biologically meaningful and appropriate for studying potamodromous Neotropical fishes. These fish might experience multiple reproductive runs, through several years which could potentially result in variable breeding outcomes, regarding the moment and place of spawning. More importantly, potential genetically distinct individuals and stocks might end up co-occurring in the same locality and time during migratory runs, or trapped between adjacent dams. This would isolate them from their original breeding grounds, leading to subsequent exogamy among different demes.

The MLE approach for admixture, which estimates population genetics parameters that maximize a likelihood function (Wang, 2003, 2022), is a powerful and scalable alternative to the more hegemonic Bayesian inference. Bayesian analysis is based on the conditional probabilities of genotypes, given some population genetics parameter and which is, in turn, highly sensible to often subjective prior probabilities (van de Schoot *et al*., 2021). Since admixture models are highly hyperdimensional (in our particular case, from 326 independent variables, for K=1, up to 6,384, for K=14), the MLE path has the advantage of high accuracy, coupled with lower computational demands (Wang, 2022), which allows for more extensive replication. Here, it showed a high degree of convergence, across replicates (many have zero variance), for different K.

We have adopted Wang’s D_LK2_ estimator for optimal K, as it was shown to be generally more reliable, over varying investigated scenarios, in comparison with the F_STIS_ estimator (Wang, 2022). The 1^st^ order MLE analysis of admixture and the DAPC both initially pointed for two groups, which did not correlate with sampling scheme. Both clusters contain members from all sampled populations. This strongly suggests the occurrence of mixed stocks of *P. lineatus*, from the main channel of the Upper Grande River. Caution was taken to dismiss panmixia as the best possibility, by directly examining the log likelihood values of alternative models, since D_LK2_ would not possibly capture K=1 (Wang, 2022), just like Evanno’s ΔK for Bayesian estimation (Janes *et al*., 2017), and given that the F_STIS_ estimator would always be zero (Wang, 2022). The results from the Bayesian analysis of admixture also reinforce the notion of several, different fish stocks in our total sample, not correlated with sampling locations or moments, supporting the hypothesis of the occurrence of mixed stocks throughout the Upper Grande River Basin.

Different values of K can reveal multiple underlying aspects of population structure and distinct stages in the population’s history (Gilbert, 2016; Janes *et al*., 2017; Gehri *et al*., 2021; Wang, 2022). The two major genetically differentiated groups defined in the 1^st^ order MLE analysis could be the reflex of existence of two main ancient breeding grounds (*e*.*g*. the Mortes, Capivari or the Aiuruoca Rivers), defining forks along migratory routes, splitting groups in the Upper Grande River. The DAPC conducted without assigning samples to localities or groups also revealed particular alleles from one of the best performing markers as being important in explaining the variation between these two DAPC defined clusters, and the cross-validation attested for its robustness. However, this method might show irregular success across particular scenarios, such as with high migration rate (as would be expected in admixed populations) and low level of differentiation among demes (as might be expected of species with high mobility), often underestimating the presence of more internal groups (Miller *et al*., 2020).

Together, these results confirm the probable existence of at least two mixed stocks along the studied area, not explained by the recent implementation of dams. Evidence for mixed stocks in *P. lineatus* were previously forwarded, for example, by Rueda *et al*. (2013), Avigliano *et al*. (2019) and indirectly by Perini *et al*. (2021). The Bayesian exploration of the admixture model suggested deeper levels of substructuring, pointing to as many as eight possible clusters in the whole sample of fish, a scenario compatible with the eight demes defined during the “peeling” of MLE layers.

Aiming to deepen the exploration of these seemingly co-occurring genetically mixed stocks of *P. lineatus* from the main channel of the Upper Grande River, the two clusters initially defined by MLE were iteratively unfolded, for two more analyses rounds. Consistent and surprisingly, every examined cluster revealed two newly characterized smaller subgroups, nested within clusters resolved in the immediate previous step (2^3^). Judging by the sorted samples of newly defined clusters from Figure 3c, the F_STIS_ estimates from Supplementary Figure 5, or the HWE analysis from MLE clusters, there would be potentially more admixture degrees to be untangled, from further hierarchical analysis, although we would reach rather critical sample sizes. The MLE defined clusters showed the expected admixture model behaviour towards higher orders. The 3^rd^ order had more genetic homogeneity within clusters and a maximization of genetic variation and distance among clusters. It also showed increasing numbers of *loci* in HWE, attesting to the meaningfulness of the hierarchical admixture analysis and the usefulness of these new microsatellite markers.

The observed stratified, self-similar, hidden pattern of the population genetic structure, across the larger sample of fish, may reveal the existence of potential different, well defined, ancestral breeding grounds, hierarchically nested within differential scale levels, throughout the basin, organized according to fractal geometry (e.g. 1^st^ order river, 2^nd^ and 3^rd^ order tributaries, and so on). If this association is confirmed, we are led to conclude, it may very well constitute the first indirect genetic evidence for a strong degree of homing behaviour for a characiformes piracema fish species. Philopatric spawning, or homing behaviour, a thoroughly investigated issue in salmonid and other migratory species, is still poorly understood in the Neotropics. It has been documented for Neotropical catfishes (*e*.*g*. Batista and Alves-Gomes, *2006; Pereira et al*., 2009). But most importantly, it was suggested for the congeneric *P. argenteus*, based on radio-tagging results from the São Francisco River Basin (Godinho and Kynard, 2006) and discussed for *P. lineatus* itself by Careaga and Carvajal-Vallejos (2019), from morphometric data. Our work revives this area of inquiry, for migratory fish in South America.

Being particularly abundant and a conspicuously competent, far-ranging migratory species, we argue that in a pre-damming past, *P. lineatus* could probably reach the innermost free-flowing tributaries for spawning, upon the colossal La Plata Basin, which cuts through a diversified range of biomes and microhabitats. We hypothesize this would likely point towards philopatric behaviour as an evolutionary stable strategy, due to the possible occurrence of distinct local adaptations, as it has been shown in other fish species (*e*.*g*. Salles *et al*., 2016; Mobley *et al*., 2019; Vøllestad and Primmer, 2019).

In conservation biology, the understanding of degrees of admixture is important in management decisions (Wang, 2003, 2022). The confirmation of mixed stocks and the hierarchical genetically stratified nature of *P. lineatus* from the main channel of an important large and intensely dammed river will likely enlighten its future research and fisheries management. If further corroborated, these findings, particularly its ensuing propositions, would have direct implications to environmental and development decision-making. It highlights the possible central relevance of key smaller tributaries (actively sought for the implementation of new small hydroelectric power plants - Ferreira *et al*. 2022), for the conservation of genetic variation, in a socio-environmentally relevant species, as it highlights the importance of free-flowing tributaries of different orders throughout the basin.

On a broader level, we raise the question whether these apparent fractal layers from the genetic structure could imply in a more widespread phenomena for piracema, other migratory fishes, or even other non-aquatic migratory organisms, showing a temporary mixture, but segregated breeding grounds. We believe this is worthy of further investigation, since fractal structures are pervasive in nature, not only in biology (*e*.*g*. Bassingthwaighte, *1992; Leggett et al*., 2019; Meyer *et al*., 2020; Sendker *et al*., 2024), but it is also conspicuously present on major geographical landscape attributes like mountain ranges, forest borders, rivers and coastlines (*e*.*g*. Xu *et al*., 1993; Andronache *et al*., 2019). The only direct discussions on the relation of fractals and genetic population structure were laid by Wingen *et al*. (2007), regarding computer simulations results on long-distance dispersal of clonal fungal spores, and by Lee (2020), exploring the fractal dimension of human genetic variability data, within and among genomes.

Our results will very likely benefit researchers and environmental managers in tackling challenging issues regarding the conservation of Neotropical migratory fishes, as they support the hypothesis of the isolating effects of dams and broodstocking over genetic variability. More importantly, the results presented here point to the potential role of tributaries of different orders in harbouring genetically diverse populations of migratory freshwater fishes.

## Acknowledgements

We thank the fishermen couple Donizete Monteiro de Paula and Sônia Bárbara de Oliveira Paula, biologists and photographers couple Lucia and André Seale, biologists João de Magalhães Lopes, Miriam Aparecida de Castro, Raquel Coelho Loures and everyone at Peixe Vivo, José Mauro Ribeiro for bioinformatics assistance, Mônica Soares da Silva for assistance with marker development, Fausto Moreira da Silva Carmo for assistance with specimen collection and DNA extraction, Ian Cosenza Irsigler for map assistance, Pedro Manoel Galetti Jr for valuable comments and Evanguedes Kalapothakis. This work was financed by Conselho Nacional de Desenvolvimento Científico e Tecnológico (CNPq grant 303023/2023-6), Companhia Energética de Minas Gerais and Agência Nacional de Energia Elétrica (CEMIG-ANEEL grant GT345), by the Coordenação de Aperfeiçoamento de Pessoal de Nível Superior - Brazil (CAPES finance Code 001), and by Fundação de Amparo à Pesquisa do Estado de Minas Gerais (FAPEMIG grant APQ-04569-10).

## Contributions

G.M.Y. conceived, conducted and supervised the study, raised funding, performed bioinformatics and population genetics analyses, wrote and edited the manuscript; G.A.S.N. performed marker development, population genetics analysis and wrote the manuscript; L.C.C. performed marker development; R.C.D.G. performed bioinformatics analysis; R.P.S. performed bioinformatics analysis; R.S.O. edited the manuscript, supervised and performed bioinformatics pipelines. All contributors read and agreed upon the final manuscript.

## Supporting Information

**Supplementary File 1:** Microsatellite DNA sequences and primers presented in the plain text format (txt) for *Prochilodus lineatus*. The order of columns is “Locus”; “Original fasta label”; “Motif”; “Repeats”; “5’-flank”; “3’-flank”; “F-primer”; “F-Tm”; “R-Primer”; “R-Tm”; “PCR product”; “Product length”; “Location”; “Scaffold or contig label in the newly presented assembly, when mapped”. This file is available at fisgshare under doi: 10.6084/m9.figshare.26848654. https://doi.org/10.6084/m9.figshare.26848654.

**Supplementary File 2:** List of DNA sequences presented in the fasta format for contigs from the original assembly from where microsatellite *loci* for *Prochilodus lineatus* were originally characterized. This file is available at fisgshare under doi: 10.6084/m9.figshare.26848015 https://doi.org/10.6084/m9.figshare.26848015.

**Supplementary File 3:** *Prochilodus lineatus* population samples, fish IDs and its rearrangements according to 1^st^, 2^nd^, and 3^rd^ order MLE defined clusters. This file is available at fisgshare under doi: 10.6084/m9.figshare.26871031. https://doi.org/10.6084/m9.figshare.26871031.

**Supplementary File 4:** Supplementary Tables, including log likelihood values of the clustering analysis for panmixia and genetic diversity estimates for *Prochilodus lineatus* MLE defined clusters from the 1^st^, 2^nd^, and 3^rd^ orders of hierarchical analysis. This file is available at fisgshare under doi: 10.6084/m9.figshare.27020857. https://doi.org/10.6084/m9.figshare.27020857.

**Supplementary File 5:** Hardy-Weinberg exact tests for MLE defined clusters of *Prochilodus lineatus*, from the 1^st^, 2^nd^, and 3^rd^ orders of hierarchical analysis. This file is available at fisgshare under doi: 10.6084/m9.figshare.27020869. https://doi.org/10.6084/m9.figshare.27020869.

**Supplementary Figures 1 – 12:** Supplementary Figures 1 through 12 are presented in a single portable document format (pdf) file, with respective captions. This file is available at figshare under doi: 10.6084/m9.figshare.26889478 https://doi.org/10.6084/m9.figshare.26889478.

